# Functional architecture of deleterious genetic variants in the Wrangel Island mammoth genome

**DOI:** 10.1101/137455

**Authors:** Erin Fry, Sun K. Kim, Sravanthi Chigurapti, Katelyn M. Mika, Aakrosh Ratan, Alexander Dammermann, Brian J. Mitchell, Webb Miller, Vincent J. Lynch

**Author notes:** These authors contributed equally (Erin Fry, Sun K. Kim, Sravanthi Chigurapti).

## Abstract

Woolly mammoths were among the most abundant cold adapted species during the Pleistocene. Their once large populations went extinct in two waves, an end-Pleistocene extinction of continental populations followed by the mid-Holocene extinction of relict populations on St. Paul Island ~5,600 years ago and Wrangel Island ~4,000 years ago. Wrangel Island mammoths experienced an episode of rapid demographic decline coincident with their isolation, leading to a small population, reduced genetic diversity, and the fixation of putatively deleterious alleles, but the functional consequences of these processes are unclear. Here we show that the Wrangel Island mammoth accumulated many putative deleterious mutations that are predicted to cause diverse behavioral and developmental defects. Resurrection and functional characterization of Wrangel Island mammoth genes carrying these substitutions identified both loss and gain of function mutations in genes associated with developmental defects (HYLS1), oligozoospermia and reduced male fertility (NKD1), diabetes (NEUROG3), and the ability to detect floral scents (OR5A1). These results suggest that Wrangel Island mammoths may have suffered adverse consequences from their reduced population sizes and isolation.

## Introduction

The end of the Pleistocene was marked by dramatic environmental change as repeated glacial cycles gave way to the warmer, more stable Holocene, including the near complete loss of the cold and dry steppe-tundra (also known as the Mammoth steppe) and the extinction of cold-adapted megafaua such as aurochs, steppe bison, cave bear, Irish elk, and woolly rhinoceros. Woolly Mammoths (*Mammuthus primigenius*) were among the most abundant cold adapted megafaunal species during the Middle to Late Pleistocene (*ca* 116-12 Kyr BP), inhabiting a large swath of steppe-tundra that extended from Western Europe, through Asia and Beringia, into North America. Paleontological and genetic data indicate that their once large populations experienced at least two demographic declines, the first during the Middle-Early Pleistocene ~285 Kyr BP (Palkopoulou et al., 2015) or Eemian interglacial ~130-116 Kyr BP (Palkopoulou et al., 2013) after which populations rebounded, and a final decline around the Pleistocene-Holocene transition (Nyström et al., 2012; 2010; Palkopoulou et al., 2013; 2015; Thomas, 2012).

Although mainland Woolly Mammoths were extinct by ~10,500 years ago, rising sea levels isolated small populations on St. Paul Island in the Bering sea ~14,000 years ago and on Wrangel Island in the Arctic sea ~9,000 years ago that survived into the Mid-Holocene. Population genetic studies have identified two distinct phases in the extinction of Woolly Mammoths (Thomas, 2012): An end-Pleistocene decline and extinction of continental populations, particularly in Northern Siberia (Nyström et al., 2012; 2010; Palkopoulou et al., 2013; 2015), followed by the mid-Holocene extinction of relict populations on St. Paul Island ~5,600 years ago (Graham et al., 2016) and Wrangel Island ~4,000 years ago (Nyström et al., 2012; 2010; Palkopoulou et al., 2013; 2015). While a combination of habitat loss and human hunting contributed to the decline and extinction of continental mammoths (Barnosky et al., 2004; Lorenzen et al., 2011; Nogués-Bravo et al., 2008), the synergistic effects of shrinking island area and freshwater scarcity caused by continued sea level rise likely caused the extinction of St. Paul Island mammoths (Graham et al., 2016).

The final causes of the extinction of Wrangel Island mammoths are unclear. However, they experienced a period of rapid demographic decline coincident with their isolation, resulting in a small population, reduced genetic diversity, recurrent breeding among distant relatives, and the fixation of deleterious alleles (Nyström et al., 2012; 2010; Pečnerová et al., 2016; Rogers and Slatkin, 2017; Thomas, 2012). These data suggest their extinction may have been associated with a ‘mutational meltdown’ (Rogers and Slatkin, 2017), but the functional consequences of putatively deleterious amino acid substitutions in the Wrangel Island mammoth are unknown. Here we identify and characterize the functional architecture of genetic variants in the Wrangel Island mammoth genome. We found that putatively damaging substitutions unique to the Wrangel Island mammoth are enriched for numerous deleterious phenotypes, such as reduced male fertility and neurological defects. Functional characterization of specific resurrected Wrangel Island mammoth genes indicates that mutations in these genes were indeed deleterious, and may have adversely effected development, reproduction, and olfaction.

## Results

To characterize the functional architecture of deleterious variants in mammoth genomes, we first identified homozygous nonsynonymous substitutions unique (private) in three extant Asian elephants (*Elephas maximus*) and three woolly mammoths, the ~44,800 year old Oimyakon mammoth (Palkopoulou et al., 2015), the ~20,000 year old M4 mammoth (Dikov, 1988; Gilbert et al., 2008; 2007; Lynch et al., 2015; Miller et al., 2008), and the ~4,300 years old Wrangel Island mammoth (Palkopoulou et al., 2015). These mammoths span the age from when mammoth populations were large and wide-spread (Oimyakon), to near the beginning of their final decline (M4), and their last known population (Wrangel Island); thus these individuals allow us to compare older variants to variants unique to the last population of mammoths. To reduce potential false positives resulting from damaged sites and other sources of error associated with ancient DNA, we hard-masked all sites that would be potentially affected by the characteristic ancient DNA patterns of cytosine deamination in single stranded overhangs from the variant-calling pipeline. This mask was applied to 10 nucleotides on both ends of the merged reads from the ancient samples. The effect of this hard masking is to reduce the total number of variant calls, but increase confidence that the called variants are real rather than artifacts of DNA preservation and damage.

Next we used PolyPhen-2 (Adzhubei et al., 2010; 2013) to computationally predict the functional impact of each private homozygous amino acid substitution (**Table 1, Fig. 1A**) and identified mouse knockout phenotypes and tissues in which these genes were enriched. We identified 115 ‘probably damaging’ amino acid variants in 112 genes in the Wrangel Island mammoth genome. These genes were enriched for 102 mouse knockout phenotypes at an FDR≤0.20 (**Supplementary Table 1**) and 63 tissues at an FDR≤0.22 (**Supplementary Table 1**) that were not observed as enriched in the deleterious variants from M4 or Oimyakon, nor any of the Asian elephants (**Fig. 1B**). Genes with ‘probably damaging’ amino acid variants in the Wrangel Island mammoth were enriched for diverse KO phenotypes, including behavioral and neurological defects such as ‘impaired righting response’ (*P*=0.008, FDR *q*=0.19), ‘abnormal brain wave pattern’ (*P*=0.017, FDR *q*=0.19), ‘catalepsy’ (*P*=0.026, FDR *q*=0.19), and ‘abnormal substantia nigra morphology’ (*P*=0.038, FDR *q*=0.20).

**Table 1.**
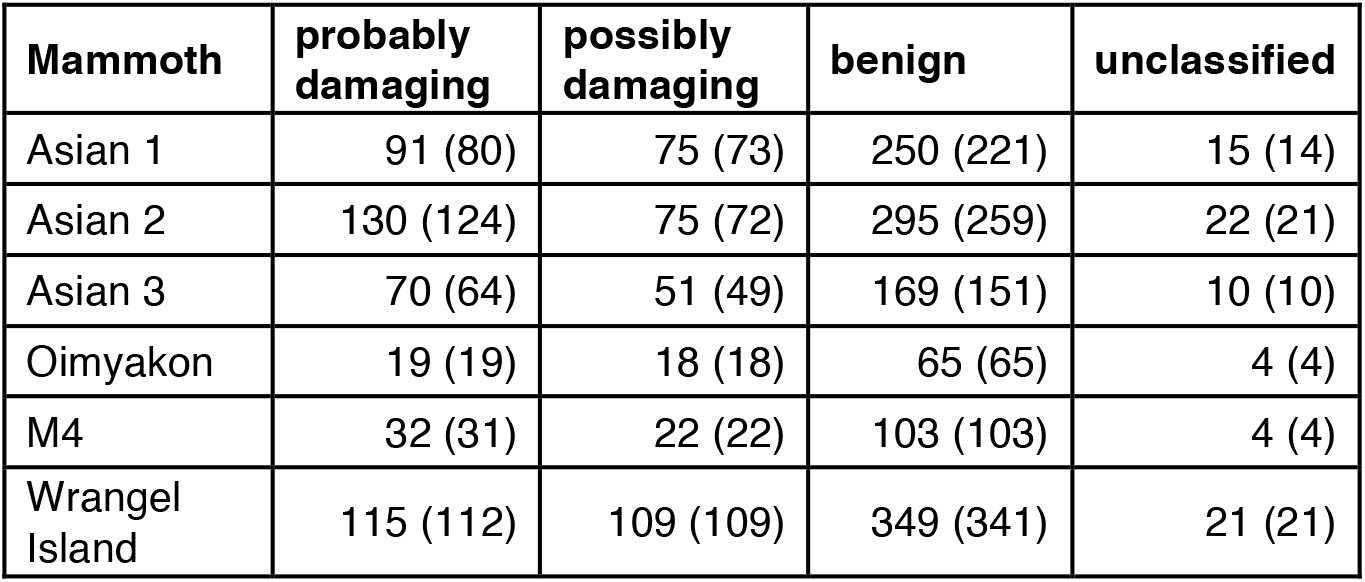
PolyPhen-2 classification of private amino acid variants in Asian elephants and woolly mammoths. (Numbers in parenthesis indicate the number of genes with those variants.)

**Figure 1.**
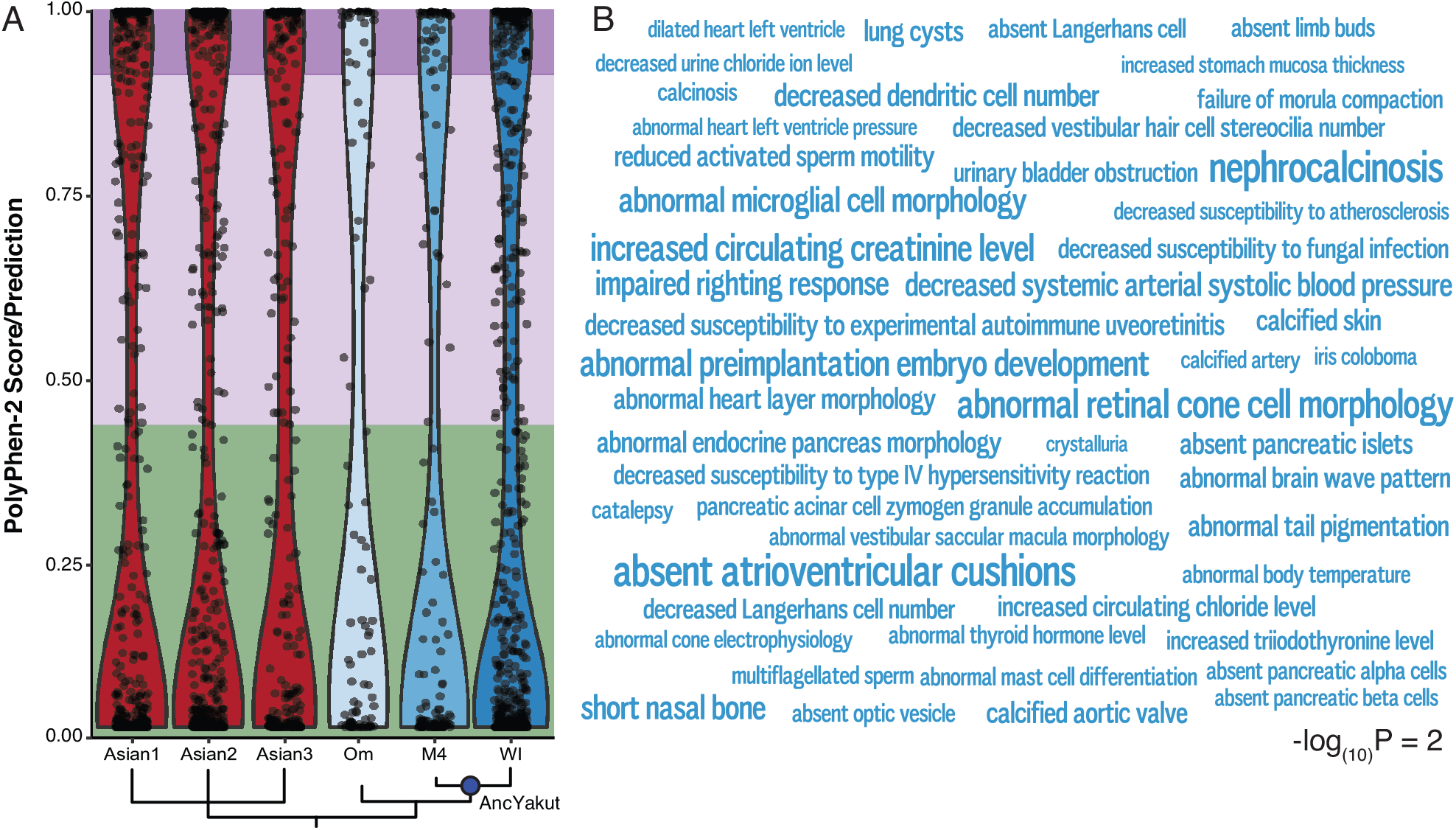
Accumulation and functional architecture of deleterious variants in the Wrangel Island mammoth genome. (**A**) Violin plot of PolyPhen-2 scores for derived, homozygous variants in each mammoth and Asian elephant. Phylogenetic relationships are shown at the bottom and the ancestral Yakut (AncYakut) node is indicated with a blue circle. Green, light purple, and dark purple backgrounds indicate variants that are predicted to be benign, possibly damaging, and probably damaging, respectively. (**B**) Word Cloud showing mouse KO phenotypes in which fixed probably damaging amino acid substitutions in the Wrangel Island mammoth are enriched. In this figure the size of each phenotype is drawn proportional to the - log_(10)_ P-value of that terms enrichment. Inset scale shows word size for phenotypes with a – log_(10)_ P-value of 2.

Among the genes with a deleterious variant with the Wrangel Island mammoth with behavioral and neurological functions is hydrolethalus syndrome protein 1 (*HYLS1*), a centriolar protein that functions in ciliogenesis (Dammermann et al., 2009). The Wrangel Island mammoth HYLS1 P119L (c.356C>T) mutation occurs in a highly conserved region of the protein, which is invariant for proline or serine across vertebrates and is therefore potentially deleterious (**Fig. 2A**). To infer if this mutation had functional consequences, we used a *Xenopus* model of ciliogenesis (Dammermann et al., 2009). Morpholino oligos (MO) targeting *Xenopus HYLS1* led to a severe defect in cilia assembly (Wilcox test, *P*=9.48×10^-6^; **Fig. 2C/H**), as previously reported (Dammermann et al., 2009). This defect was rescued by addition of MO-resistant wild-type *Xenopus HYLS1* (Wilcox test, *P*=1.71×10^-4^; **Fig. 2D/H**), but not a variant incorporating the equivalent P119L mutation into *Xenopus* HYLS1 (S186L) (Wilcox test, *P*=2.56×10^-5^; **Fig. 2E/H**). The HYLS1 S186L mutant did, however, appropriately localize to centrioles and did not have any dominant negative effects in the absence of depletion of the endogenous protein (**Fig. 2F/G**). Mutations in HYLS1 underlie hydrolethalus syndrome (HLS [MIM: 236680]), a perinatal lethal developmental disorder characterized by severe brain malformation including hydrocephalus and absent midline structures (Mee et al., 2005), as well as Joubert syndrome (JBTS [MIM: 213300]), a milder disorder characterized by defects in the cerebellum and brain stem leading to impaired balance and coordination (Oka et al., 2016), suggesting the HYLS1 P119L mutation may have had adverse developmental consequences.

**Figure 2.**
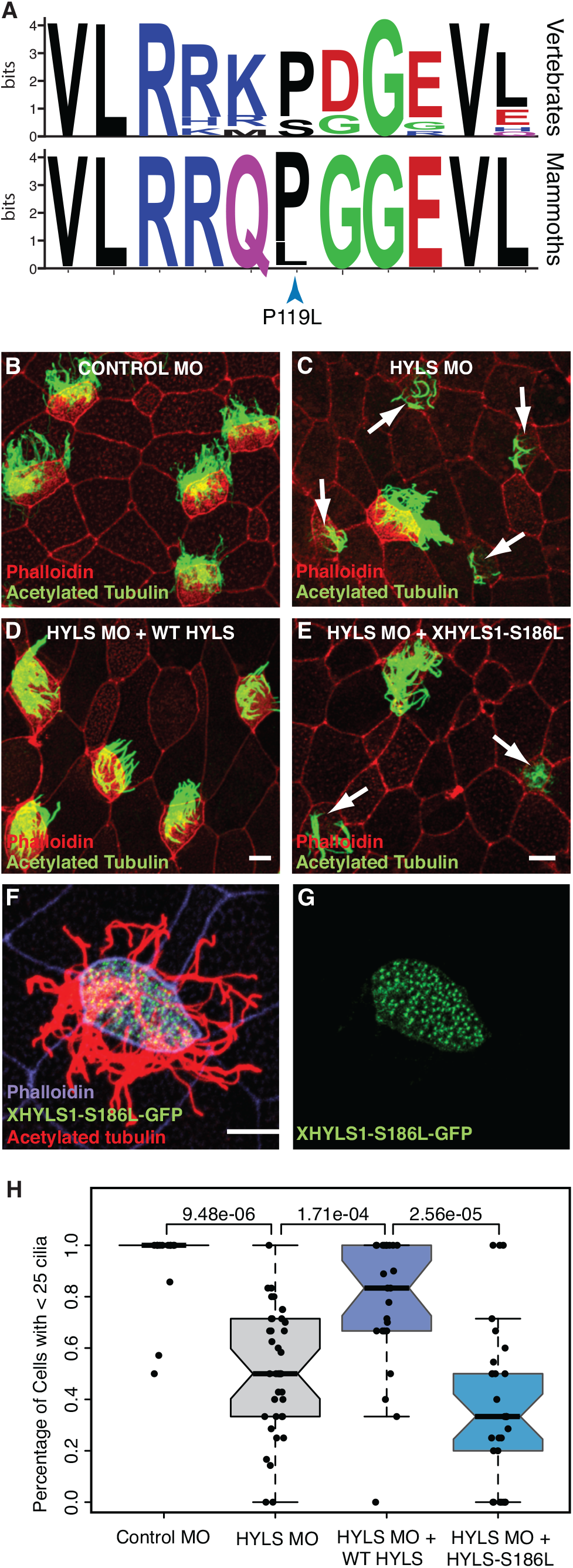
The Wrangel Island mammoth HYLS1 P119L mutant is dysfunctional. (**A**) Sequence logo showing conservation of HYLS1 around the Wrangel Island mammoth P119L mutation. Upper, logo from 100 vertebrates. Lower, logo from mammoths (Oimyakon, M4, and Wrangel Island). (**B-E**) Representative images of *Xenopus* multiciliated cells from embryos injected with Control MO (**B**), HYLS1 MO (**C**), HYLS1 MO and WT XHYLS1 (**D**) and HYLS1 MO and XHYLS1-S186L (**E**) stained with acetylated tubulin to mark cilia (green) and phalloidin to mark cell boundaries (red), note: arrows identify cells with defective ciliogenesis (scale bars = 10μm). (**F-G**) Multiciliated cell from a *Xenopus* embryo injected with mRNA encoding XHYLS1-S186L-GFP shows proper localization to the centrioles (green) and proper formation of cilia marked with acetylated tubulin (red) with cell boundaries marked with phalloidin (purple)(scale bar = 10μm). (**H**) Quantification of ciliogenesis defect scoring cells with > 25 cilia (Wilcox test *P*-values are shown).

Defects in sperm morphology are among the most common consequences of reduced genetic diversity and inbreeding (Asa et al., 2007; Shorter et al., 2017), and several knockout phenotypes unique to the Wrangel Island mammoths are related to sperm biology such as ‘reduced activated sperm motility’ (*P*=0.011, FDR *q*=0.19), ‘multiflagellated sperm’ (*P*=0.023, FDR *q*=0.19) and ‘abnormal sperm axoneme morphology’ (*P*=0.045, FDR *q*=0.19); ‘sperm flagellum’ was also the tissue most significantly enriched deleterious variants (*P*=0.001, FDR *q*=0.22). Among the genes with a ‘probably damaging’ amino acid substitution in the Wrangel Island mammoth associated with sperm defects and male infertility is *Naked cuticle 1* (*NKD1*), which encodes a passive antagonist of the Wnt/TCF-LEF signaling pathway (Angonin and Van Raay, 2013; Van Raay et al., 2007; 2011). The Wrangel Island mammoth A88V substitution occurred at a site that is nearly invariant for alanine or serine in diverse vertebrates (**Fig. 3A**), suggesting it may have had functional consequences.

**Figure 3.**
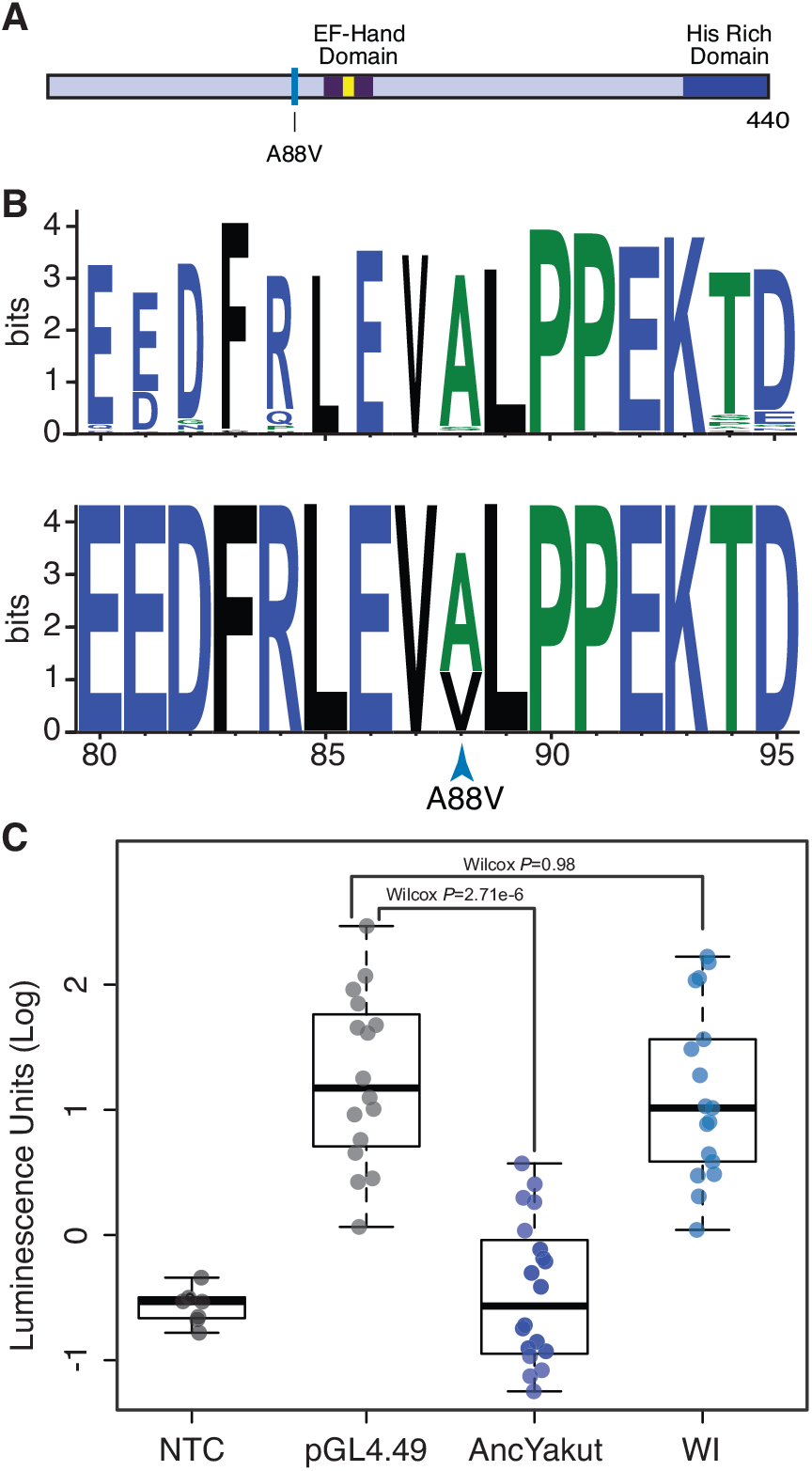
The Wrangel Island mammoth NKD1 A88V substitution is a loss of function mutation. (**A**) The Wrangel Island mammoth A88V mutation is located near the EF-hand domain (dark blue), the calcium binding motif is shown in yellow. (**B**) Sequence logo showing conservation of NKD1 around the Wrangel Island A88V mutation. Upper, logo from 100 vertebrates. Lower, logo from mammoths (Oimyakon, M4, and Wrangel Island). (**C**) Luciferase expression from the pGL4.49[*luc2P*/TCF-LEF/Hygro reporter vector in elephant fibroblasts transfected with control vector, AncYakut *NKD1*, or Wrangel Island (WI) Mammoth *NKD1*, and treated with the WNT signaling agonist CHIR99021. Background luminescence of non-transfected cells (NTC).

To determine if the NKD1 A88V (c.163C>T) substitution had functional effects, we resurrected the Wrangel Island/M4 ancestral (AncYakut, **Fig. 3A**) and Wrangel Island *NKD1* genes and tested their ability to antagonize Wnt-signaling in elephant dermal fibroblasts transiently transfected with a luciferase reporter vector containing a minimal promoter and eight copies of a TCF-LEF response element (pGL4.49[*luc2P*/TCF-LEF/Hygro]) and treated with a small molecule agonist of the Wnt-signaling pathway (CHIR99021). The AncYakut NKD1 reduced luminescence to background levels in response to CHIR99021 treatment (Wilcox test, *P*=2.71×10^-6^). In stark contrast however, the Wrangel Island NKD1 did not affect luciferase expression (Wilcox *P*=0.98), indicating the NKD1 A88V substitution is a loss of function mutation (**Fig. 3B**). Transgenic mice with loss of function mutations in NKD1 have dysregulated Wnt/beta-catenin signaling in the testis leading to abnormal seminiferous tubule morphology, small seminiferous tubules, small testis, oligozoospermia, and reduced fertility (Li et al., 2005; Zhang et al., 2007), suggesting this substitution may have affected male fertility in Wrangel Island mammoths.

Although deleterious Wrangel Island mammoth variants are enriched in diverse KO phenotypes, many are related to the pancreas such as ‘abnormal endocrine pancreas morphology’ (*P*=0.011, FDR *q*=0.19), ‘absent pancreatic islets’ (*P*=0.013, FDR *q*=0.19), ‘pancreatic acinar cell zymogen granule accumulation’ (*P*=0.019, FDR *q*=0.19), ‘absent pancreatic alpha cells’ (*P*=0.023, FDR *q*=0.19), and ‘absent pancreatic beta cells’ (*P*=0.027, FDR *q*=0.19). Among the genes with deleterious Wrangel Island variants annotated with ‘abnormal endocrine pancreas morphology’ is *NEUROGENIN 3* (*NEUROG3*), which encodes a basic helix-loop-helix transcription factor that is required for endocrine cell development. The ‘probably damaging’ NEUROG3 G195E (c.584G>A) substitution in the Wrangel Island mammoth NEUROG3 protein occurred at a site that is nearly invariant for glycine across mammals (**Fig. 4A/B**) within an LXXLL motif in the C-terminal transcriptional activation domain (Smith et al., 2004), suggesting it may alter protein function.

**Figure 4.**
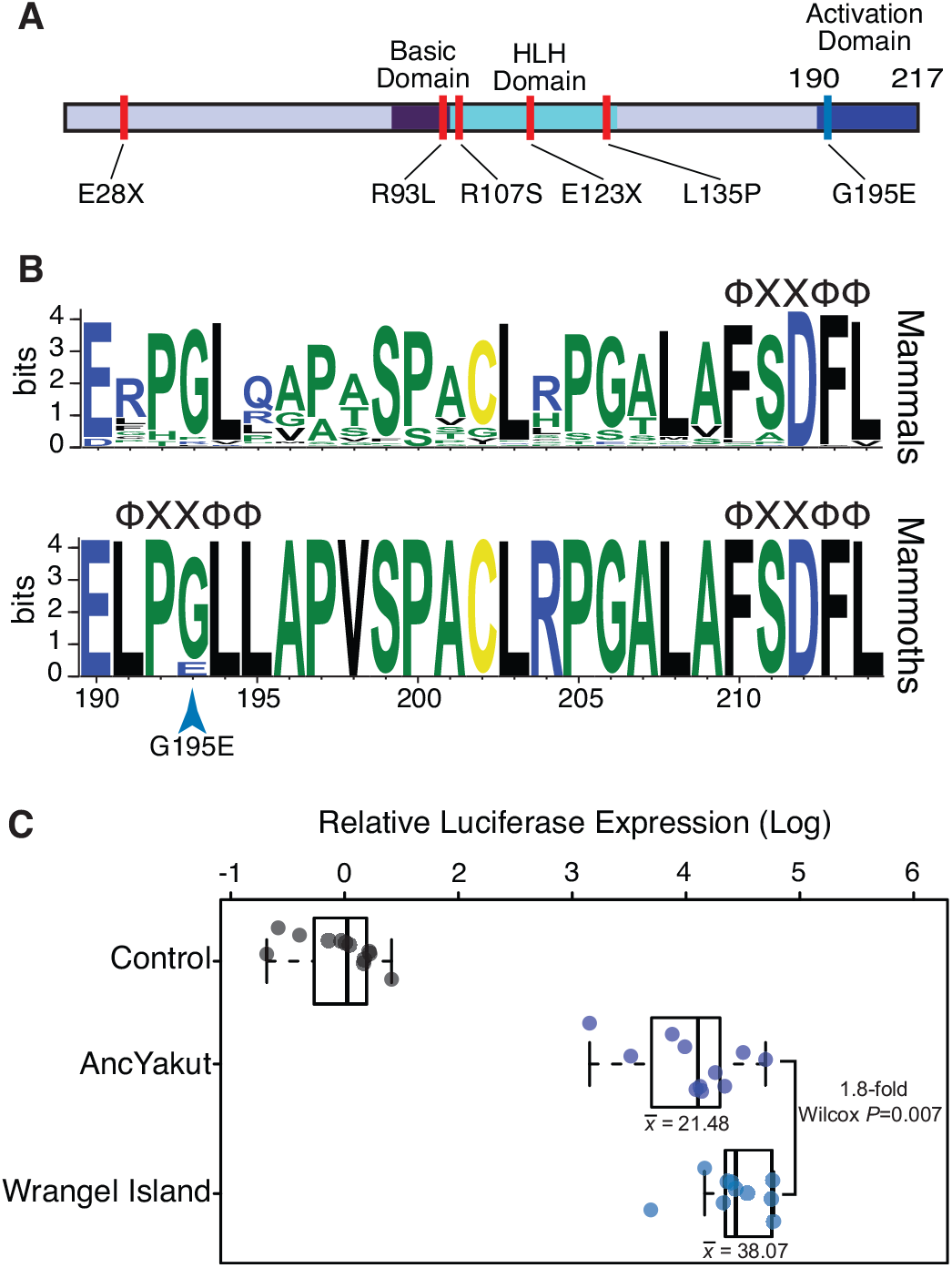
The Wrangel Island mammoth NEUROG3 is a hypermorph. (**A**) The Wrangel Island mammoth G195E mutation is located in the C-terminal transactivation domain of NEUROG3, location of mutations associated with human disease are shown in red. (**B**) Sequence logo showing conservation of NEUROG3 transactivation domain in mammals (upper) and mammoths (lower; Oimyakon, M4, and Wrangel Island). The location of ΦXXΦΦ co-factor interaction motifs are shown (Φ, any hydrophobic amino acid; X, any amino acid). (**C**) Relative luciferase expression from the pGL3 [*luc/6x-PAX4E/minP*] reporter vector in elephant fibroblasts transfected with control vector, AncYakut *NEUROG3*, or Wrangel Island Mammoth *NEUROG3*.

To determine if the G195E substitution had functional effects, we resurrected the AncYakut and Wrangel Island *NEUROG3* genes and tested their ability to transactivate luciferase expression from a reporter vector containing a minimal promoter and six repeats of the *PAX4* E-box (pGL3[*luc/6x-PAX4E/minP*]) in transiently transfected elephant dermal fibroblasts (**Fig. 4C**). The Wrangel Island NEUROG3 transactivated luciferase expression from the pGL3[*luc/6x-PAX4E/minP*] reporter vector more strongly than the AncYakut NEUROG3 protein (1.8-fold, Wilcox *P*=0.007), indicating the NEUROG3 G195E substitution is a hypermorphic mutation. Loss of function mutations in the human *NEUROG3* gene cause congenital malabsorptive diarrhea (DIAR4 [MIM:610370]), a disorder characterized by neonatal diabetes, chronic unremitting malabsorptive diarrhea, vomiting, dehydration, and severe hyperchloremic metabolic acidosis (Pinney et al., 2011; Rubio-Cabezas et al., 2011; Wang et al., 2006). *NEUROG3* knock-out mice die postnatally from diabetes (Rubio-Cabezas et al., 2011) suggesting the NEUROG3 G195E substitution may have affected insulin signaling in Wrangel Island Mammoths.

A previous study of the Wrangel Island mammoth genome found a high rate of pseudogenization in olfactory receptors (Rogers and Slatkin, 2017), which have greatly expanded in the elephant lineage (Niimura et al., 2014) and generally evolve rapidly through both adaptive and neutral birth-death processes (Nei et al., 2008). Consistent with these data, we found that ordorant receptors were the largest class of gene (21/115) with ‘probably damaging’ mutations in the Wrangel Island mammoth. Among the olfactory receptors with ‘probably damaging’ amino acid substitution is *OR5A1*, which encodes the mammalian β-ionone sensor (Jaeger et al., 2013). β-ionones are of a family of closely related aroma compounds known as rose ketones, which are the major contributor to the aroma of flowers such roses and violets. Remarkably a human D183N polymorphism (rs6591536), close to the Wrangel Island mammoth OR5A1 S193F (c.578C>T) substitution (**Fig. 5A**), underlies differential sensitivity to β-ionone in humans (Jaeger et al., 2013). β-ionone sensitive individuals, for example, can more easily distinguish food and beverages with added β-ionone than insensitive individuals, and typically describe β-ionone as "fragrant" and "floral" whereas insensitive individuals describe β-ionone as smelling like “sour/acid/vinegar’’and “sharp/pungent/acid”.

**Figure 5.**
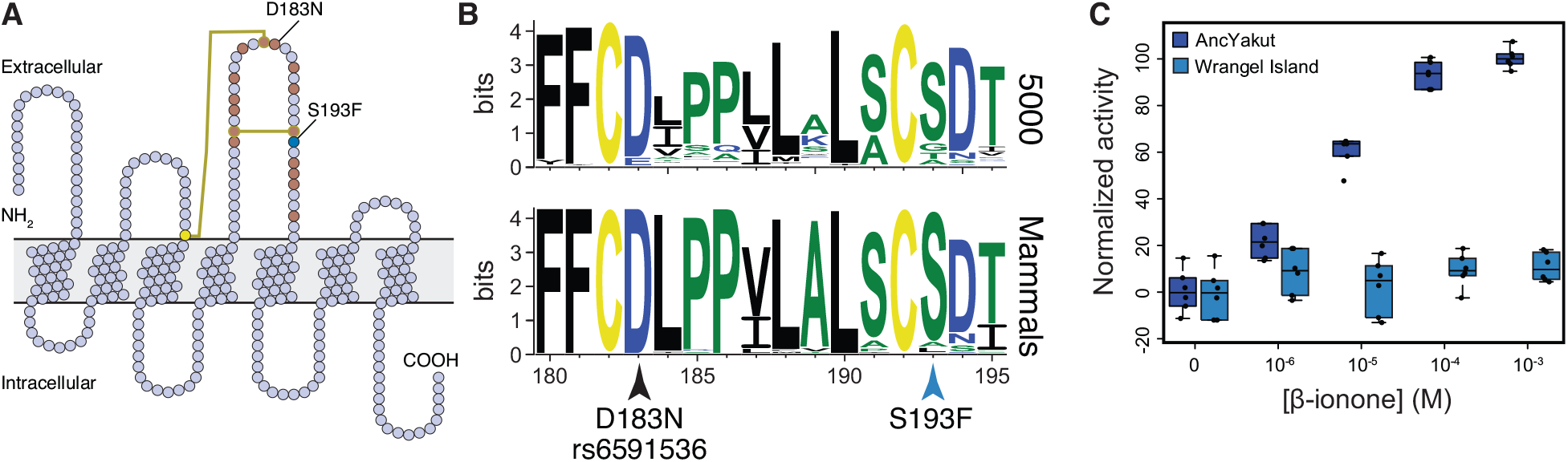
The Wrangel Island mammoth β-ionone sensor OR5A1 is non-functional. (**A**) Snake diagram of the Wrangel Island mammoth OR5A1 protein. The location of cysteine residues and disulfide bonds are shown as yellow circles and lines, respectively. The location of residues previously shown by high throughput mutagenesis to affect receptor function are shown as brown circles (Mainland et al., 2014). The location of the Wrangel Island mammoth S193F substitution and the human D183N polymorphism are also shown. (**B**) Sequence logo showing conservation of OR5A1 AAs 180-195 from 5000 randomly selected odorant receptors (upper) and mammals (lower). Location of the S193F substitution and the D183N polymorphism are shown. (C) Dose response curve showing normalized activity of the AncYakut and Wrangel Island mammoth OR5A1 odorant receptors to β-ionone. Data shown are standardized to non-transfected Hana3a cells and no β-ionone, *n*=6

Both the human D183N and Wrangel Island mammoth OR5A1 S193F substitutions occur in a disulfide bonded extracellular loop that plays a role in ligand recognition (**Fig. 5A**) (Mainland et al., 2014; Man et al., 2004; Yu et al., 2015; Zhuang et al., 2009). This site is also nearly invariant for serine in mammalian OR5A1 orthologs as well as 5000 diverse olfactory receptors paralogs (**Fig. 5B**) suggesting the S193F substitution may affect receptor function. To determine if the S193F substitution had functional consequences, we resurrected the AncYakut and Wrangel Island mammoth *OR5A1* genes and tested their sensitivity to β-ionone using the Hana3A odorant receptor assay (Saito et al., 2004; Zhuang and Matsunami, 2008). We found that the AncYakut OR5A1 was strongly activated by β-ionone. In stark contrast the Wrangel Island mammoth OR5A1 completely lacked β-ionone sensitivity, indicating that the OR5A1 S193F substitution is a loss of function mutation (**Fig. 5C**). Forbs were prominent in the diet of late Quaternary megafauna (Willerslev et al., 2014), including mammoths, suggesting the S193F substitution in OR5A1 may have altered the ability of Wrangel Island mammoths to detect one of their major food sources.

## Discussion

The final causes of the extinction of Wrangel Island mammoths are mysterious, but it is clear that Wrangel Island mammoths experienced an episode of demographic decline coincident with their isolation leading to a chronically small population. The minimum effective population size to prevent the loss genetic diversity in wild populations has been estimated to be ~500, while the minimum effective population size to prevent the accumulation of deleterious mutations is ~1000 (Nunney and Campbell, 1993; Thomas, 1990). The effective population size of Wrangel Island mammoths has been estimated to be ~300-500 (Palkopoulou et al., 2015) (Nyström et al., 2012; 2010), close to the 150-800 individual carrying capacity of the island (Nyström et al., 2012). These data suggest that the Wrangel Island mammoth population was too small to effectively purge deleterious mutations. Consistent with expectations for small populations, previous studies of Wrangel Island mammoths have found signatures of reduced genetic diversity, recurrent breeding among distant relatives, and the fixation of putatively deleterious alleles (Nyström et al., 2012; 2010; Palkopoulou et al., 2015; Pečnerová et al., 2016; Rogers and Slatkin, 2017; Thomas, 2012). These data suggest that deleterious mutations accumulated in Wrangel Island mammoths in response to long term low effective population sizes, and may have contributed to their extinction (Rogers and Slatkin, 2017) (Palkopoulou et al., 2015).

We found that the Wrangel Island mammoth genome had numerous fixed substitutions that are predicted to be deleterious and that these substitutions were enriched for specific abnormal phenotypes compared to older, continental populations of mammoths, and Asian elephants. Consistent with our computational analyses of putatively deleterious substitutions, we validated gain or loss of function mutations in *HYLS1*, *NKD1*, *NEUROG3*, and *OR5A1*, confirming that at least some predicted deleterious mutations were indeed function altering. The loss of function mutation in OR5A1, for example, likely altered the ability of Wrangel Island mammoths to detect β-ionone and thus floral scents whereas the loss of function mutation in NKD1 may have affected male fertility in Wrangel Island mammoths. The loss of function mutation in HYLS1 may have had more widespread affects given the widespread importance of cilia in vertebrate development (Badano et al., 2006). In contrast to the other genes we tested, the Wrangel Island specific mutation in *NEUROG3* is a hypermorph rather than a loss of function suggesting it may have caused gain of function phenotypes related to the development and function of pancreatic beta cells (Smith et al., 2004).

Unfortunately while mammoths are an excellent case study for the evolution of derived phenotypes (Lynch et al., 2015) and the genomics of isolation and extinction (Palkopoulou et al., 2015; Rogers and Slatkin, 2017), we are unable to do the kinds of forward and reverse genetic experiments that generally establish causal associations between genotypes and phenotypes. Thus, this study has obvious limitations. We infer, for example, the functional consequences of amino acid substitutions using a computational model that compares the properties amino acids and the likelihood of observing the derived substitution given the pattern of amino-acid variation at that site in orthologous genes (Adzhubei et al., 2010; 2013). We cannot know, however, whether the potentially deleterious effects of amino acid substitutions are buffered through epistasis and suppressed. Similarly we infer the phenotypic consequences of deleterious amino acid variants by reference to mouse knockout and human disease data, assuming that the same gene has the same role across species. While this is often the case, development is plastic and the gene regulatory, morphogenetic, and structural bases of homologous characters can diverge through a process of developmental systems drift (Liao and Zhang, 2008; Lynch, 2009; True and Haag, 2001; Wang and Sommer, 2011). Even with these limitations, our computational inferences and functional validation provide strong evidence that Wrangel Island mammoths likely suffered adverse consequences from their prolonged isolation and small population size, providing a cautionary tale for modern conservation biology.

## Methods

### Genome assembly

Details of the sequencing protocol for the Oimyakon and Wrangel Island mammoths can be found in Palkopoulou et al. (2015) and for the Asian elephants, M25, and M4 in Lynch et al. (2015). Sequences from the Asian elephants were aligned to the reference genome from the African Elephant (loxAfr3) using the Burrows Wheeler Aligner (Li and Durbin, 2010) with default parameters (BWA version 0.5.9-r16). The reads were subsequently realigned around putative indels using the GATK (DePristo et al., 2011) IndelRealigner (version 1.5-21-g979a84a), and putative PCR duplicates were flagged using the MarkDuplicates tool from the Picard suite (version 1.96).

The sequences from the mammoths were treated separately to account for DNA damage in the sequences. Putative adapter sequences were removed and we merged overlapping paired-end reads using available scripts (Kircher, 2012). We required an overlap of at least 11 nucleotides between the mates, and only pairs that could be merged were retained for subsequent analyses. The merged reads were aligned to the genome from the African elephant (loxAfr3) using BWA with default parameters, and only the mapped reads that were longer than 20 bps were retained for the subsequent SNP calls. The reads were realigned using the GATK IndelRealigner and putative PCR duplicates were flagged using MarkDuplicates, similar to the process described for the modern genomes. We also limited the incorporation of damaged sites into the variant-calling pipeline by hard-masking all sites that would be potentially affected by the characteristic ancient DNA patterns of cytosine deamination in single stranded overhangs. This mask was applied to 10 nucleotides on both ends of the merged reads from the ancient samples.

Single-nucleotide variants (SNVs), i.e. positions in the African elephant reference assembly at which we detected a nucleotide different from the reference in at least one of the Asian elephant or mammoth individuals were identified using SAMtools (Li et al., 2009) (version 0.1.19), which was applied with “-C50” to adjust the mapping quality of the reads with multiple mismatches. We did not call differences in regions where the reference base was unknown, and the calls were limited to regions that were covered at least 4 times, and at most 350 times by the sequences in these samples. We then identified homozygous nonsynonymous substitutions unique (private) to each elephant and mammoth genome; these homozygous nonsynonymous substitutions were used in downstream analyses.

The M25 genome is likely a composite of multiple mammoth individuals (Rogers and Slatkin, 2017), therefore, we did not include M25 in our functional analyses (described below). However, because we are primarily interested in private nonsynonymous variants within each elephant and mammoth genome we were able to take advantage of the composite nature of the M25 genome by excluding homozygous nonsynonymous substitutions identified in the three Asian elephants and the three mammoths that were also observed in M25. The rationale for this filtering process is that any homozygous nonsynonymous substitution observed in either the Asian elephants or the three mammoths that is also observed in M25 is not truly a private variant. We thus identified 106 fixed amino acid substitutions in 99 genes in the Oimyakon mammoth, 162 fixed amino acid substitutions in 143 genes in the M4 mammoth, and 594 fixed amino acid substitutions in 525 genes in the Wrangel Island mammoth.

### Functional annotation

To infer the putative functional consequences of amino acid substitutions in the Wrangel Island mammoth genome, we focused on fixed (homozygous) derived variants that were not observed in the other mammoth or elephant samples. Our focus on fixed, derived variants excludes possible deleterious heterozygous variants but reduces the risk of mis-classifying amino acid variants that arise from DNA damage. We used PolyPhen-2 to classify amino acid substitutions as ‘benign’, ‘possibly damaging’, or ‘probably damaging’ (Adzhubei et al., 2010; 2013) and Enrichr (Chen et al., 2013; Kuleshov et al., 2016) to infer the functional consequences of fixed ‘probably damaging’ amino acid substitutions in each mammoth and elephant. We then intersected these phenotypes to identify those unique to the Wrangel Island mammoth. We report (unique) enriched mouse knockout phenotypes at an FDR≤0.20. We also used the same approach to identify tissues in which genes with ‘probably damaging’ amino acid substitutions in the Wrangel Island mammoth are enriched.

### Data availability

Homozygous nonsynonymous substitutions unique (private) in the three extant Asian elephants and mammoths and PolyPhen-2 functional annotations for each variant are available at Galaxy and are included as supplementary materials.

### Selection of target genes for functional validation

We manually curated each gene with a predicted probably damaging amino acid substitutions in the Wrangel Island mammoth based on literature searches and selected targets for functional validation based on three criteria: 1) The Wrangel Island mammoth specific amino acid substitution must have been classified by PolyPhen-2 as ‘probably damaging’ with a pph2_prob score ≥0.958; 2) Prior studies (based on literature reviews) must have identified the molecular function for that gene; and 3) The ability to design straightforward experimental systems in which to test the function of ancestral and derived amino acid variants. Finally, we selected genes for functional validation that had high read coverage to ensure base calls were correct, and excluded substitutions at CpG sites. Based on these criteria we selected HYLS1, NKD1, NEUROG3, and OR5A1 for functional valiation.

### HYLS1 functional validation

*Xenopus* embryos were acquired by in vitro fertilization using standard protocols (Sive et al., 2000) approved by the Northwestern University Institutional Animal Care and User Committee. Previously validated MOs (GeneTools) were used (Control MO, 5’-CCTCTTACCTCAGTTACAATTTATA-3’; HYLS-1.1,5’-GAACTGCCTGTCTCGAAGTGACATG-3’;XHYLS-1.2,5’-GAACTGCCTGTCTCTCAGTGACATG-3’ (Dammermann 2009). Full length XHYLS1 and the Wrangel mammoth mutant *Xenopus* equivalent XHYLS1-S186L were cloned into pCS2+ and fused with GFP at the N terminus. mRNA of the pCS2 constructs was prepared using in vitro transcription with SP6 (Promega). Morpholinos and mRNA were coinjected into each blastomere at the 2-4 cell stage using a total of 50-75 ng of morpholino and 500pg-1ng mRNA per embryo. Embryos were allowed to develop until stage 28 then fixed with 4%PFA in PBS for 2 hrs at RT. For antibody staining embryos were blocked for 1 hr in PBS with 0.1% Triton and 10% Normal Goat Serum prior to overnight incubation with primary antibody (Acetylated tubulin, Sigma T6793). Fluorescent secondary Abs (Jackson Labs) were incubated overnight after a full day of washing in PBS-0.1 %Triton. After secondary washing embryos were stained with fluorescently tagged phalloidin to mark the cell boundaries. Imaging was performed on a laser-scanning confocal microscope (A1R; Nikon) using a 60× oil Plan-Apo objective with a 1.4 NA.

### NKD1 functional validation

To infer if the A88V substitution had functional affects, we synthesized the ancestral mammoth (AncYakut, A88) and Wrangel Island (V88) *NKD1* genes with mouse codon usage and cloned each gene into the mammalian expression vector pcDNA3.1+C-DYK. Next we tested their ability to antagonize luciferase expression from the pGL4.49[*luc2P*/TCF-LEF/Hygro luciferase reporter vector, which drives luciferase expression from a minimal promoter and eight copies of a TCF-LEF response element upon activation of Wnt-signaling. African elephant primary dermal fibroblasts (San Diego Zoo, “Frozen Zoo”) were grown at 37°C/5% CO2 in a culture medium consisting of FGM/MEM (1:1) supplemented with 10% FBS, 1% Glutamine, and 1% penstrep. Confluent cells in 96-well plates in 60 μL of Opti-MEM (GIBCO) were transfected with 100 ng of the luciferase reporter plasmid pGL4.49[*luc2P*/TCF-LEF/Hygro, 100 ng of the AncYakut or Wrangel Island Mammoth NKD1 expression vector, and 10 ng of pRL-null with 0.1μL of PLUS reagent (Invitrogen) and 0.3 μL of Lipofectamine LTX (Invitogen) in 20 μL of Opti-MEM. The cells were incubated in the transfection mixture for 6 hr until the transfection media was replaced with culture media and supplemented with the small molecule Wnt-signalig agonist CHIR99021. 48 hours after transfection, Dual Luciferase Reporter Assays (Promega) began with incubating the cells for 15 min in 20 μL of 1x passive lysis buffer. Luciferase and Renilla activity was measured with the Glomax multi+detection system (Promega). We standardized luciferase activity values to Renilla activity values and background activity values by measuring luminescence in wells lacking the NKD1 expression vector.

### NEUROG3 functional validation

To determine if the G195E substitution had functional effects, we synthesized the ancestral mammoth (AncYakut, G195) and Wrangel Island (E195) *NEUROG3* genes with mouse codon usage and cloned each gene into the mammalian expression vector pcDNA3.1+C-DYK. Next we tested their ability to transactivate luciferase expression from the pGL3 luciferase reporter vector containing a minimal promoter and six repeats of the *PAX4* E-box (pGL3 [*luc/6x-PAX4E/minP*]); the *PAX4* E-box has previously been shown to physically bind NEUROG3 and drive luciferase expression in reporter assays (Smith et al., 2004). African bush elephant primary dermal fibroblasts (San Diego Zoo, “Frozen Zoo”) were grown at 37°C/5% CO2 in a culture medium consisting of FGM/MEM (1:1) supplemented with 10% FBS, 1% Glutamine, and 1% penstrep. Confluent cells in 96-well plates in 60 μL of Opti-MEM (GIBCO) were transfected with 100 ng of the luciferase reporter plasmid 6x-PAX4E_pGL3-Basic, 100 ng of the AncYakut or Wrangel Island Mammoth NEUROG3 expression vector, and 10 ng of pRL-null with 0.1μL of PLUS reagent (Invitrogen) and 0.3 μL of Lipofectamine LTX (Invitogen) in 20 μL of Opti-MEM. The cells were incubated in the transfection mixture for 6 hr until the transfection media was replaced with culture media. 48 hours after transfection, Dual Luciferase Reporter Assays (Promega) began with incubating the cells for 15 min in 20 μL of 1x passive lysis buffer. Luciferase and Renilla activity was measured with the Glomax multi+detection system (Promega). We standardized luciferase activity values to Renilla activity values and background activity values by measuring luminescence in wells lacking the NEUROG3 expression vector.

### OR5A1 functional validation

We used the Hana3A odorant receptor assay to determine if the Wrangel Island mammoth OR5A1 S193F substitution had functional effects. This assay system utilizes HEK293 cells stably expressing accessory proteins required for signal transduction by odorant receptors, including RTP1L, RTP2, REEP1, and G_αolf_ (Hana3A cells) (Saito et al., 2004; Zhuang and Matsunami, 2008). We synthesized the ancestral mammoth (AncYakut, S193) and Wrangel Island (F193) *OR5A1* genes with human codon usage and cloned each gene into the mammalian expression vector pcDNA3.1+C-DYK. Next we followed the Hana3A odorant receptor assay protocol to assay their sensitivity to β-ionone (Zhuang and Matsunami, 2008) with slight modifications (described next). Hana3A cells were maintained at 37°C/5% CO2 in a culture medium consisting of FGM/MEM (1:1) supplemented with 10% FBS, 1% Glutamine, and 1% penstrep. Confluent cells in 96-well plates in 60 μL of Opti-MEM (GIBCO) were transfected with 100 ng of the cAMP responsive CRE-Luc luciferase reporter plasmid, 100 ng of the AncYakut or Wrangel Island Mammoth *OR5A1* expression vector, and 10 ng of the SV40-Renilla with 0.1μL of PLUS reagent (Invitrogen) and 0.3 μL of Lipofectamine LTX (Invitogen) in 20 μL of Opti-MEM. The cells were incubated in the transfection mixture for 6 hr until the transfection media was replaced with culture media. 48 hours after transfection, media was replaced with serum-free CD293 suspension culture medium (GIBCO) supplemented with L-glutamine and increasing concentrations of β-ionone (Sigma). Relative luminescence was assayed four hours after treatment with β-ionone using the dual Luciferase Reporter Assays (Promega) as described above. We standardized luciferase activity values to Renilla activity values and background activity values by measuring luminescence in wells lacking the *OR5A1* expression vector.

## AUTHOR CONTRIBUTIONS

E.F. performed NEUROG3 functional experiments and co-wrote the manuscript, S.K.K performed HYLS1 functional experiments and A. D. and B. J. M. co-wrote the manuscript, S. C. performed the NKD1 and OR5A12 functional experiments, K. M. M. curated variants, A.R. performed genome analyses, W. M. performed genome analyses and co-wrote the manuscript, V. J. L. performed variant analyses and co-wrote the manuscript.

## ACKNOWLEDGEMENTS

We thank H. Matsunami for the Hana3a cell line. This research was supported by a National Institutes of Health–National Institute of General Medical Sciences grant to B.J. M. (R01GM089970)

